# Unsaturated fatty acid alters the immune response in non-small cell lung adenocarcinoma through regulation of HMGB1 trafficking

**DOI:** 10.1101/2023.11.08.566231

**Authors:** Breanna Cole-Skinner, Nicole M. Andre, Zachary Blankenheim, Kate Root, Glenn E. Simmons

**Affiliations:** Department of Molecular Microbiology and Immunology, University of Missouri, Columbia; Department of Biomedical Sciences, College of Veterinary Medicine, Cornell University, Ithaca; Department of Biomedical Sciences, University of Minnesota School of Medicine, Duluth

## Abstract

Cancer cell evasion of the immune response is critical to cancer development and metastases. The ability of clinicians to kickstart the immune system to target these rogue cells is an ever-growing area of research and medicine. In this study, we delved into the relationship between lipid metabolism, High Mobility Group Box 1 protein (HMGB1), and immune regulation within non-small cell lung adenocarcinoma (NSCLC), shedding light on novel therapeutic avenues and potential personalized approaches for patients. We found that the expression of stearoyl CoA desaturase 1 (SCD1) was decreased in NSCLC tumors compared to normal tissues. This emphasized the critical role of lipid metabolism in tumor progression. Interestingly, monounsaturated fatty acid (MUFA) availability impacted the expression of programmed death receptor ligand −1 (PD-L1), a pivotal immune checkpoint target in lung cancer cells and immune cells, suggesting a novel approach to modulating the immune response. This study uncovered a complex interplay between HMGB1, SCD1, and PD-L1, influencing the immunological sensitivity of tumors. Our work underscores the importance of understanding the intricate relationships between lipid metabolism and immune modulation to develop more effective NSCLC treatments and personalized therapies. As we continue to explore these connections, we hope to contribute to the ever-evolving field of cancer research, improving patient outcomes and advancing precision medicine in NSCLC.

## Introduction

Non-small cell lung adenocarcinoma (NSCLC) is still a significant global health concern. As immunotherapeutic treatment options such as immune checkpoint blockade (ICB) become more common, many patients do not benefit from these advancements. As many as 80% of patients recommended for ICB do not respond or become refractory to treatment [1–4]. Unfortunately, the mechanism behind ICB treatment failure is unknown. This highlights a pressing need for a deeper understanding of the molecular mechanisms governing lung adenocarcinoma progression and its immune microenvironment.

Several studies have underscored the pivotal role of aberrant metabolism in shaping the behavior of cancer cells, with a particular focus on the influence of lipid metabolism [5–9]. This emerging body of evidence compels us to explore how lipid metabolic alterations impact tumor development and immune responses within the tumor microenvironment of lung cancer.

Earlier studies have shown that High Mobility Group Box 1 protein (HMGB1) drives the activity of the pro-inflammatory transcription factor NF-kB, a pivotal player in the immune response [10–12]. Although HMGB1 participates in the inflammatory response, the release of HMGB1 has also been shown to induce immunosuppression in tumors [13,14]. Interestingly, inhibition of HMGB1 improved therapeutic response [14]. The secretion of HMGB1 is regulated by lysine acetylation, which masks its nuclear localization sequence, allowing for nuclear-cytoplasmic transport and later release [15–17]. Sirtuin deacetylases remove these acetyl groups from HMGB1, preventing release in an unsaturated lipid-dependent manner, illustrating the relationship between HMGB1 and fatty acid metabolism [18–23]. As both fatty acid synthesis and HMGB1 participate in cancer development, we embarked on investigating the mechanistic underpinnings of the relationship between lipid metabolism, immune regulation, and its potential impact on cancer therapy.

In the current study, we examined The Cancer Genome Atlas (TCGA, n=542) and the Clinical Proteomic Tumor Analysis Consortium (CPTAC, n=115) Lung adenocarcinoma (LUAD) datasets to view the relationship between lipid metabolism and HMGB1 in NSCLC tumors [24]. Previous studies have reported that the presence of unsaturated fatty acids affects the expression of proteins involved in tumor growth [25]. We focused our analysis on specific proteins in lipogenic and HMGB1/RAGE pathways. Our findings revealed a relationship between stearoyl CoA desaturase (SCD1) and HMGB1 protein in the tumors of NSCLC patients. Our *in vitro* studies illustrated that altering SCD1 activity led to changes in HMGB1 release and concomitant changes in the expression of PD-L1 on the surface of lung cancer cells and innate immune cells. Preliminary examination of a small NSCLC patient population suggested a relationship exists between serum HMGB1 and tumor-associated PD-L1.

Overall, our research unveils an uncharted axis involving MUFA, HMGB1, and PD-L1 in NSCLC, shedding light on the intricate lipid metabolism network, immune regulation, and potential therapeutic targets within this challenging disease. These findings show the potential promise of targeting lipid metabolism in precision medicine and developing innovative immunotherapeutic approaches to treat NSCLC.

## Results

### Expression of stearoyl CoA desaturase is decreased in non-small lung adenocarcinoma tumors

Patients with lung cancer have been shown to have altered metabolism in critical pathways [26]. Earlier studies by our lab and others have shown that alterations in lipid metabolism are often essential to how cancer cells behave [5,7,27–29]. We were interested in investigating how lipid metabolism is altered within the lung tumors of patients. To address this question, we analyzed the transcriptome from the lung cancer patients (TCGA LUAD, n=542). Our earlier experience with stearoyl desaturase also led us to integrate this into the lung cancer cohort [29]. Within this patient population, SCD1 gene expression was significantly decreased in lung tumors compared to normal adjacent tissue (Figure 1A).

**Figure 1.**
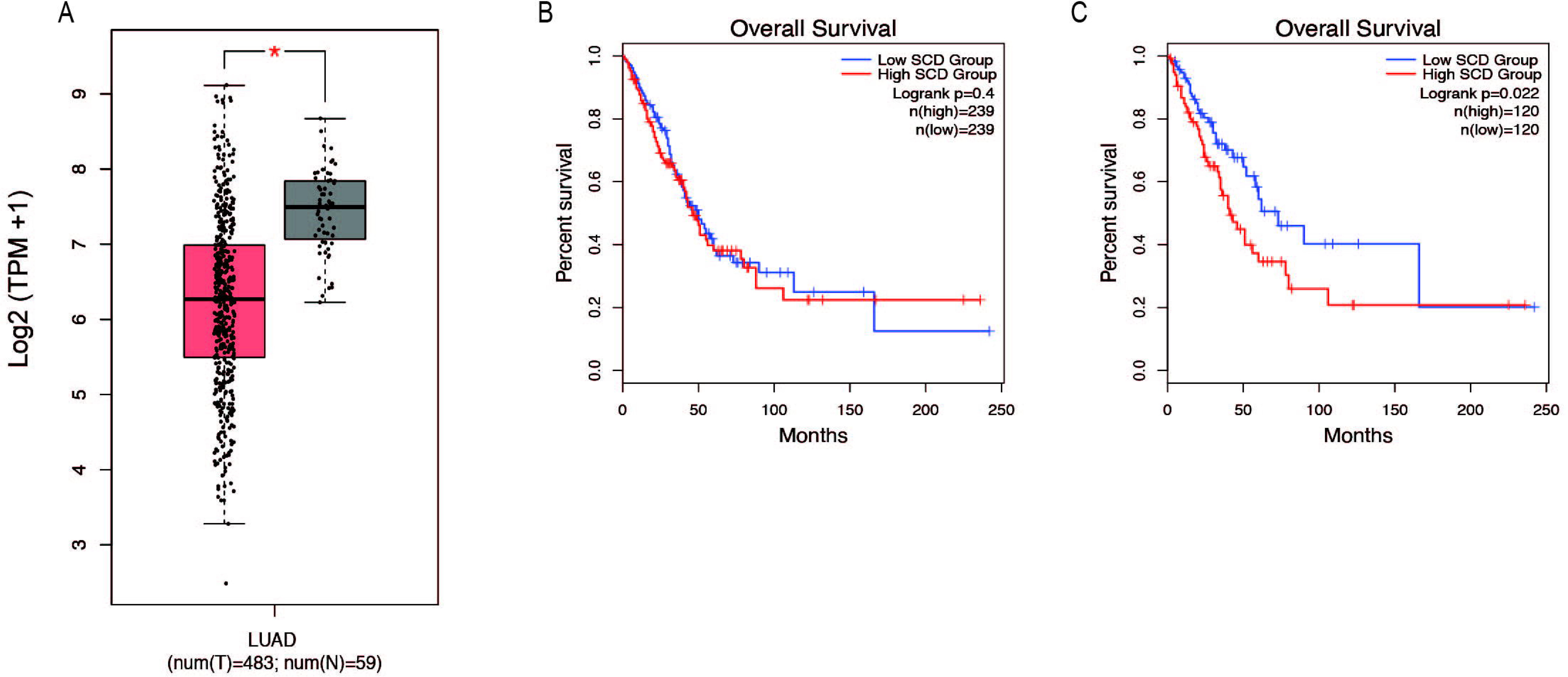
SCD1 gene expression is decreased in NSCLC tumors and associated with survival. A) Analysis of TCGA lung adenocarcinoma RNA-seq datasets from UCSC Xena project comparing SCD1 gene expression in lung tumors vs normal tissue using GEPIA webserver. B) Kaplan-Meier plot of median overall survival (OS) in groups with either high or low expression of SCD1. analysis based on gene expression. C) Quartile overall survival in groups with either high or low SCD1 gene expression. For differential gene expression, method for differential analysis is one-way ANOVA, using disease state (Tumor vs Normal) as variable for calculating differential expression. For Survival plots, a Log-rank test was done to determine significance of the difference in survival between each group.

Interestingly, the impact of SCD1 gene expression on survival was only clear in the upper and lower quartiles of the population, suggesting an outlier effect may be involved (Figure 1B and 1C). Still, seeing such a significant difference in SCD1 expression between normal and malignant tissues suggests that the activity of SCD1 and de novo production of monounsaturated fatty acids (MUFA) takes part in the development of lung tumors.

### Monounsaturated fatty acid promotes differential expression of proteins in lung cancer cells

The production of unsaturated fatty acids is essential for supporting healthy cells. However, during carcinogenesis increases in cellular metabolism, especially glycolysis, increases fatty acid production (Pouyafar et al., 2019). Fatty acid synthesis, by default produces saturated fat. The accumulation of saturated fat is toxic to cells and requires conversion to an unsaturated fatty acid via stearoyl CoA desaturases (SCD1 and SCD5). Therefore, increased unsaturated fatty acid can result from the increases in glycolysis and lipogenesis. To determine the effect of increased unsaturated fat on lung cancer proteome, we briefly deprived lung adenocarcinoma cells (A549) of exogenous fatty acids using SCD1 inhibitor (A9395762) and delipidated serum, followed by acute exposure to saturated, mono, or poly-unsaturated fatty acids. We quantified the cellular proteome using tandem mass tags, comparing relative expressions across the BSA, Oleate-BSA, Arachidonate-BSA, or Palmitate-BSA treated samples. Approximately 1,000 proteins were enriched in unsaturated (mono and poly) fatty acid-treated samples compared to BSA control (saturated fatty acids had the opposite effect on many of these proteins) (Figure 2A). Gene ontology showed that many enriched genes were associated with metabolic pathways and cellular response to infection (Figure 2B). Within the differentially enriched and repressed protein, one High mobility group box 1 protein (HMGB1) was associated with the immune response and poor prognosis in cancer patients (Figure 2C). Based on this, we investigated the relationship between HMGB1 and lipid metabolism in lung cancer.

**Figure 2.**
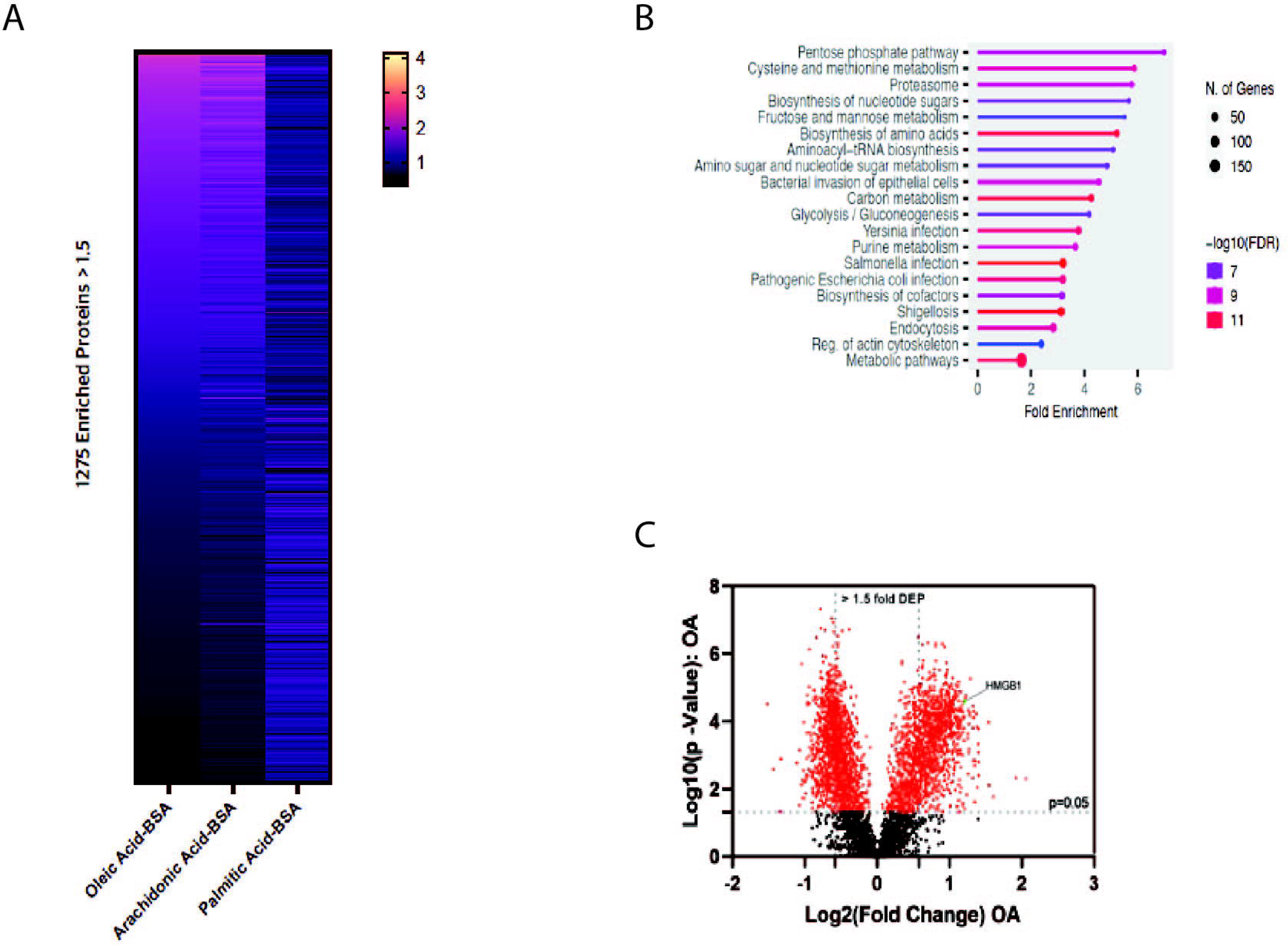
MUFA increases expression of a subset of proteins in NSCLC. A) Overview of proteomic analysis of A549 lung cancer cells following 16-hour delipidation and 4-hour fatty acid replenishment with either oleate-BSA, arachidonate-BSA, or palmitate-BSA. B) Gene Ontology of unsaturated fatty acid-enriched peptides in A549 Cells using ShinyGO **(REF).** C) Volcano plot of differential expressed peptides in A549 lung cancer cells following delipidation and oleate-BSA replenishment.

### HMGB1 and SCD1 genes are inversely related in lung cancer patients

It has been reported that HMGB1 protein plays a role in cancer progression and response to therapy. Based on earlier reports, we wanted to determine the relationship between HMGB1 and the production of unsaturated fatty acids in tumors. To explore this question, we accessed the TCGA LUAD dataset and plotted the expression of the HMGB1 gene in tumors and normal tissue. There was trivial difference in the level of HMGB1; however, there was a much larger distribution of expression in lung tumor groups (Figure 3A). Interestingly, patients with elevated HMGB1 had worse survival outcomes compared to the HMGB1 low group (Figure 3B, 3C). To confirm the presence of a relationship between HMGB1 and SCD1, we measure the correlation between the expression of both transcripts within the LUAD cohort data. A small but significant positive correlation existed between SCD1 and HMGB1 RNA in patients’ tumors (Figure 3D). This seemed counter to our working hypothesis, so we next analyzed the proteome of lung cancer tumors using the CPTAC Apollo dataset. We chose a subset of proteins involved in lipogenesis and lipid packaging, along with HMGB1 and SCD1. HMGB1 protein was inversely related to many lipogenic and lipid packaging proteins, including SCD1 and sterol response element binding factor 1 (SREBF1) (Figure 3E). These results suggest that in tumors, the level of unsaturated fatty acids has a regulatory role in the expression of HMGB1 protein.

**Figure 3.**
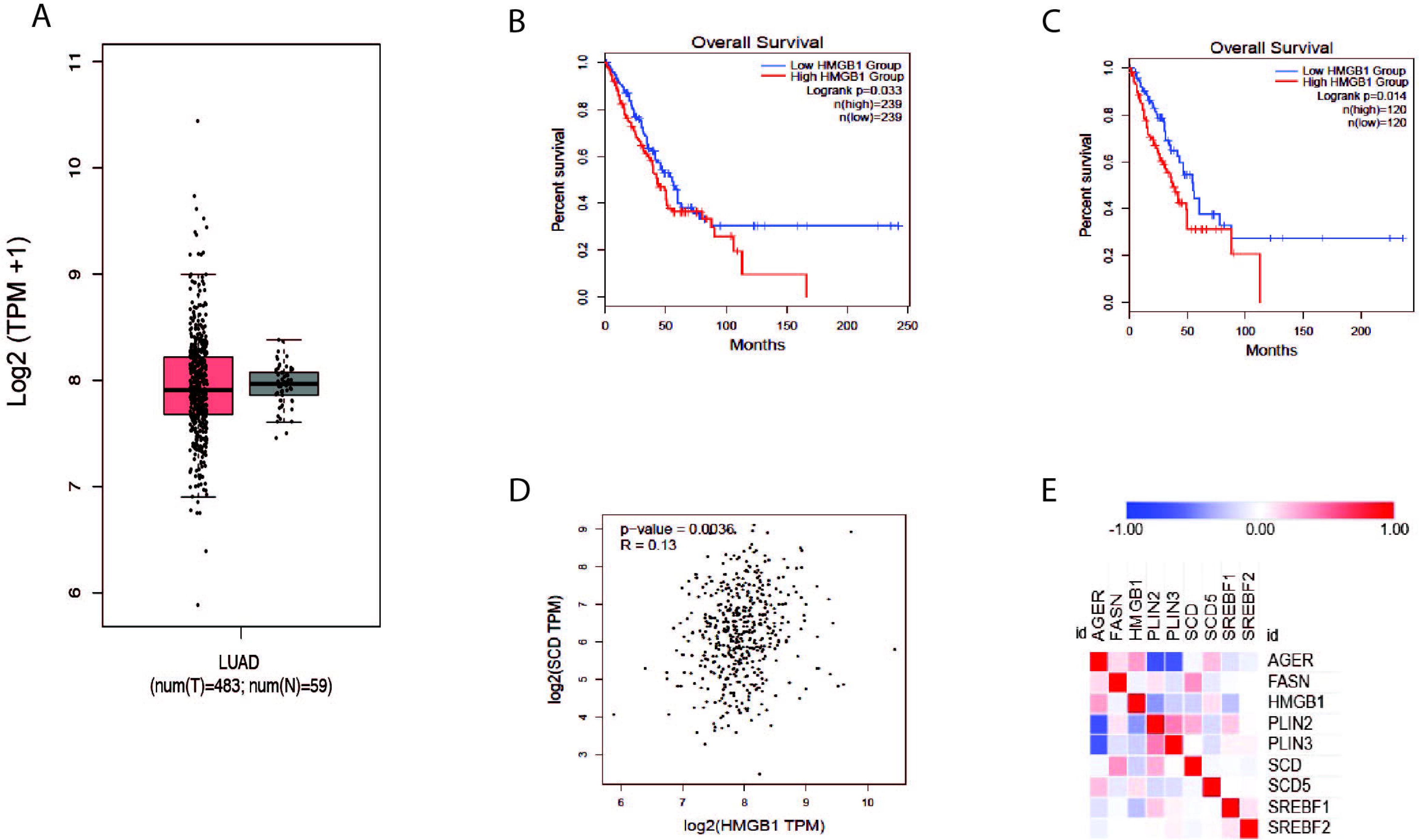
Proteins involved in HMGB1 and MUFA synthetic pathways are inversely correlated in NSCLC. A) Analysis of TCGA lung adenocarcinoma RNA-seq datasets from UCSC Xena project comparing HMGB1 gene expression in lung tumors vs normal tissue using GEPIA webserver. B) Kaplan-Meier plot of median overall survival (OS) in groups with either high or low expression of HMGB1. C) Quartile overall survival in groups with either high or low HMGB1 gene expression. D) Pearson correlation analysis between the expression of SCD1 and HMGB1 in lung cancer patients. E) Heatmap generated from Clinical Proteomic Tumor Analysis Consortium lung adenocarcinoma dataset comparing the expression proteins in lipogenic (FASN, SREBF1, SREBF2, SCD, SCD5, PLIN2, and PLIN3) and HMGB1 (AGER and HMGB1) pathways.

### Inhibition of SCD1 promotes the release of HMGB1 protein from cultured lung cancer cells

Many studies have shown that SCD1 is vital for rapid cell growth. In some instances, SCD1 has been proposed as a target for anti-cancer therapy [31–34]. Earlier work from our lab has shown that SCD1 expression is correlated with survival in patients with clear cell renal cell carcinoma (ccRCC) [35]. Based on this, we looked to determine how HMGB1 was affected by the inhibition of SCD1. Treating cells with low micromolar amounts of the SCD1 inhibitor did not affect cell viability or HMGB1 RNA levels but significantly decreased SCD1 mRNA (Figure 4A, B). Further, increased concentrations of the SCD1 inhibitor, A9395762, led to a dose-dependent decrease in intracellular HMGB1 and a concomitant increase in extracellular HMGB1 (Figure 4C, D). Thus, these data show that SCD1 activity does not affect the transcription of the HMGB1 gene; instead, it regulates the localization of HMGB1 within lung cancer cells.

**Figure 4.**
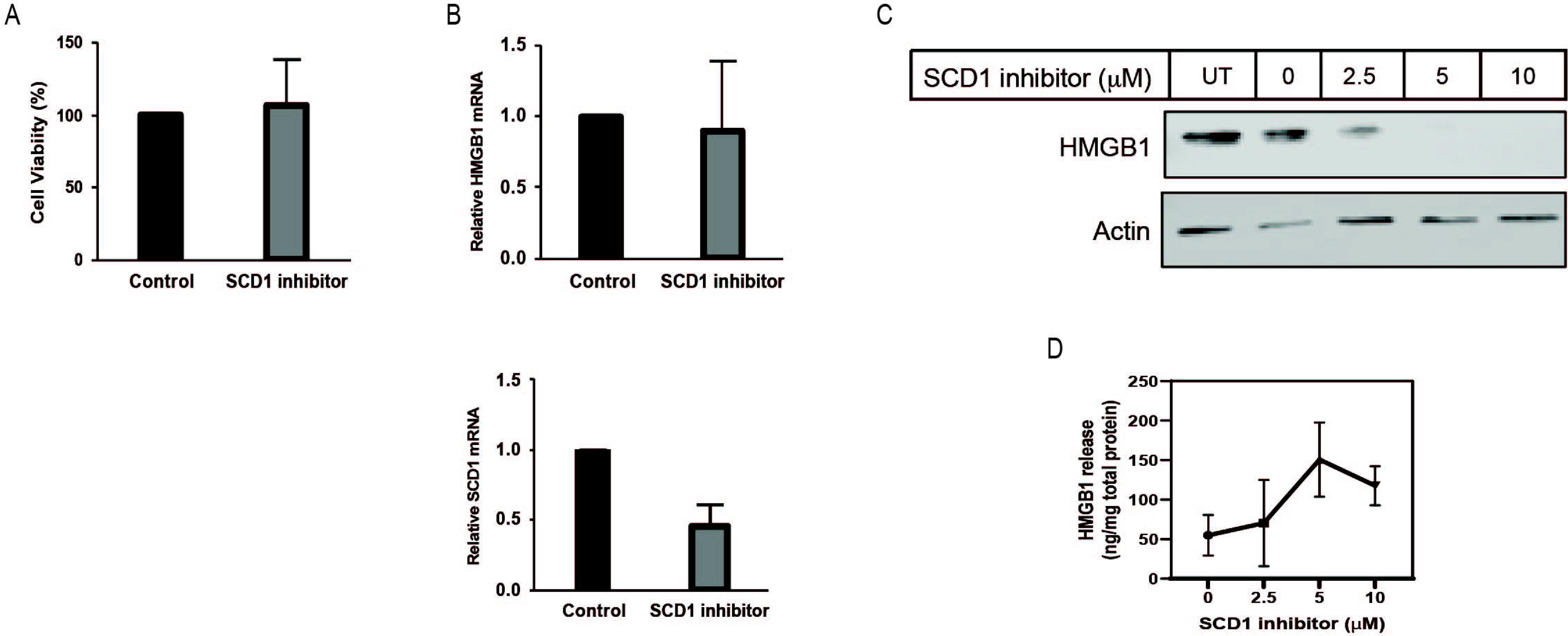
SCD inhibition promotes release of HMGB1 from lung cancer cells. A) Cell-titer Glo viability assay of cells treated for 24 hours with 2µM SCD1 inhibitor, A939572. B) Real-time quantitative PCR analysis for HMGB1 and SCD1 mRNA in A549 cells treated 24 hours with 2µM SCD1 inhibitor. C) Immunoblot analysis of HMGB1 expression in A549 cells treated 24 hours with increasing concentration of SCD1 inhibitor. D) HMGB1-specific enzyme-linked immunosorbent assay on extracellular media from A549 cells treated 24 hours with increasing concentration of SCD1 inhibitor. Error bars represent the standard error of the mean (S.E.M.) of 3 independent experiments. Blots are representative images of three independent experiments.

### Monounsaturated fatty acid increases the retention of HMGB1 in lung cancer cells

Since the inhibition of SCD1 appears to have such an impact on HMGB1 release, we directly investigated the effect of MUFA on HMGB1. Using multiple lung cancer cell lines (A549, HCC827, H1299, and H23), we measured the level of intracellular HMGB1 following our delipidation procedure. When cells were deprived of lipids, HMGB1 release was increased, but the re-introduction of MUFA significantly suppressed HMGB1 release in all cell lines evaluated (Figure 5A, B). To demonstrate how lipid supplementation affected HMGB1 within cells, we transfected GFP-tagged HMGB1 into A549 cells and exposed cells to delipidation and lipid replenishment. Besides the increased molecular weight, we confirmed that HMBG1-GFP behaved similarly to endogenous HMGB1 in lung cancer cells (Figure 5D). Once transfected, HMGB1-GFP was enriched in the cytoplasm of cells supplemented with MUFA following delipidation (Figure 5C, E). This suggests that MUFA is either blocking late steps in nuclear-cytoplasm-extracellular secretion or that MUFA is promoting the re-uptake of secreted HMGB1.

**Figure 5.**
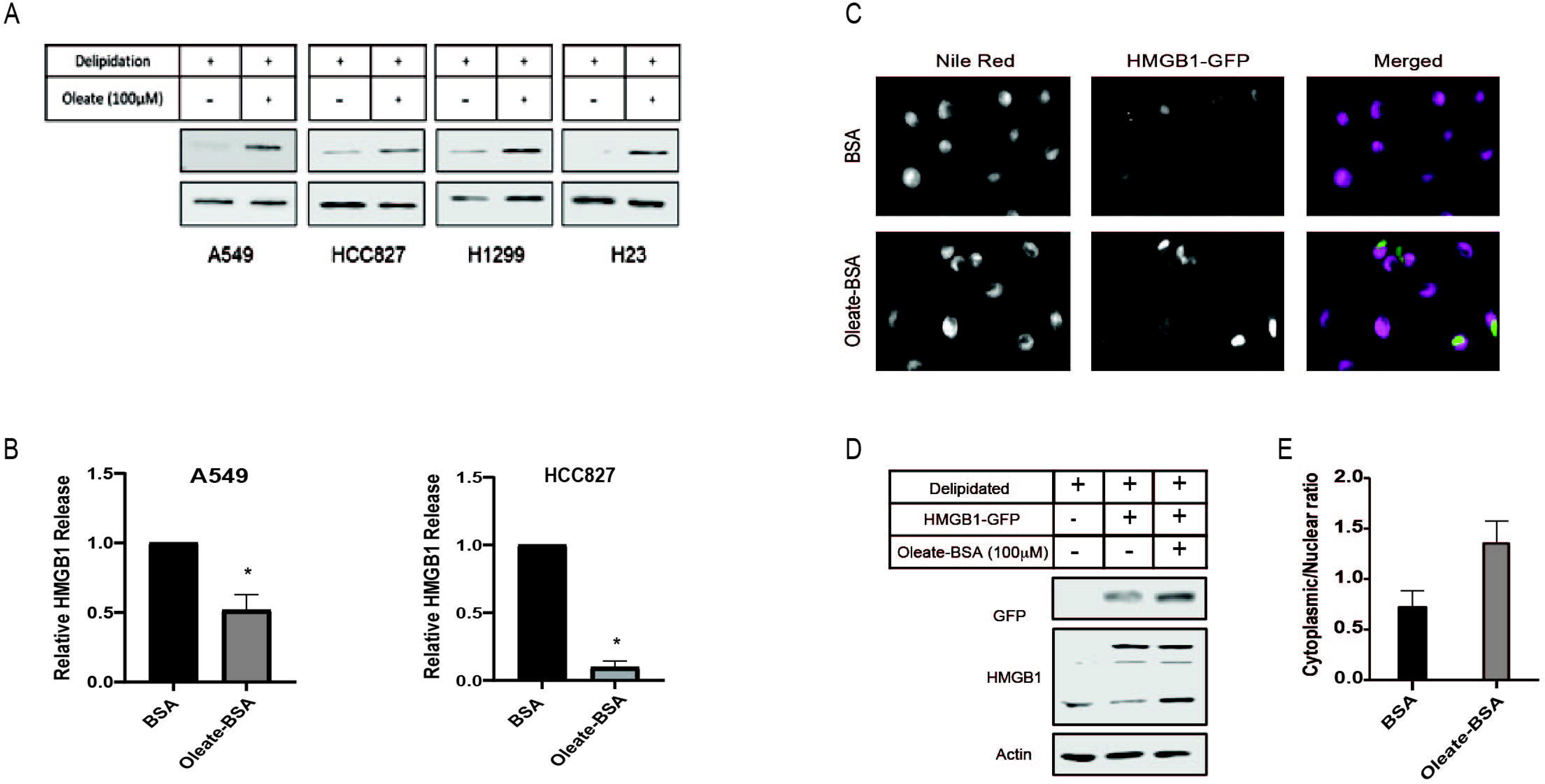
MUFA increases retention of HMGB1 in lung cancer cells. A) Immunoblot protein analysis of lung cancer cells lines treated 16 hours with 1µM SCD1 inhibitor and 4-hours replenished with oleate-BSA. B) HMGB1-specific ELISA on extracellular media from A549 and HCC827 lung cancer cells following 16-hour delipidation and 4-hour oleate-BSA replenishment. C) Fluorescence microscopy of HMGB1-GFP transfected A549 cells following 16-hour delipidation and 4-hour oleate-BSA replenishment and neutral lipid staining with Nile Red. D) Immunoblot protein analysis of HMGB1-GFP transfected A549 cells from Figure 5C. E) Quantitative analysis of GFP signal localization in transfected A549 cells in Figure 5C. Error bars represent the standard error of the mean (S.E.M.) of 5 independent experiments. Blots are representative images of three independent experiments.

### MUFA and SIRT1 cooperate to retain HMGB1 inside lung cancer cells

HMGB1 has been proven to move throughout the cell due to post-translational modifications. Several labs have shown acetylation to regulate the ability of HMGB1 to leave the nucleus and exit the cell [36–39]. Acetylation of HMGB1 is conducted by a host of acetyltransferases [40]. Conversely, removal of acetyl groups from HMGB1 is conducted by histone deacetylases (HDACs), and in particular one family of HDAC proteins known as sirtuins. Evidence suggests that SIRT1 removes acetyl groups from HMGB1 in a lipid dependent manner [21,41]. To determine the role of deacetylases in MUFA-dependent regulation of HMGB1 release, we treated A549 cells with sirtuin inhibitor, cambinol, and measured the intracellular and extracellular HMGB1 levels. In the presence of the inhibitor, we noted a decrease in cell associated HMGB1 and a concomitant increase in HMGB1 release (Figure 6A, B). Sirtuin inhibition appeared to stimulate HMGB1 release, but to demonstrate that MUFA was involved, we delipidated cells in the presence and absence of cambinol, followed by brief exposure to MUFA. Following pretreatment with sirtuin inhibitors, the application of MUFA led to a decrease in HMGB1 release and an increase in intracellular protein (Figure 6C, D). Together, these data are evidence that the relationship between HMGB1 and sirtuin is responsible for the retention of HMGB1 in lung cancer.

**Figure 6.**
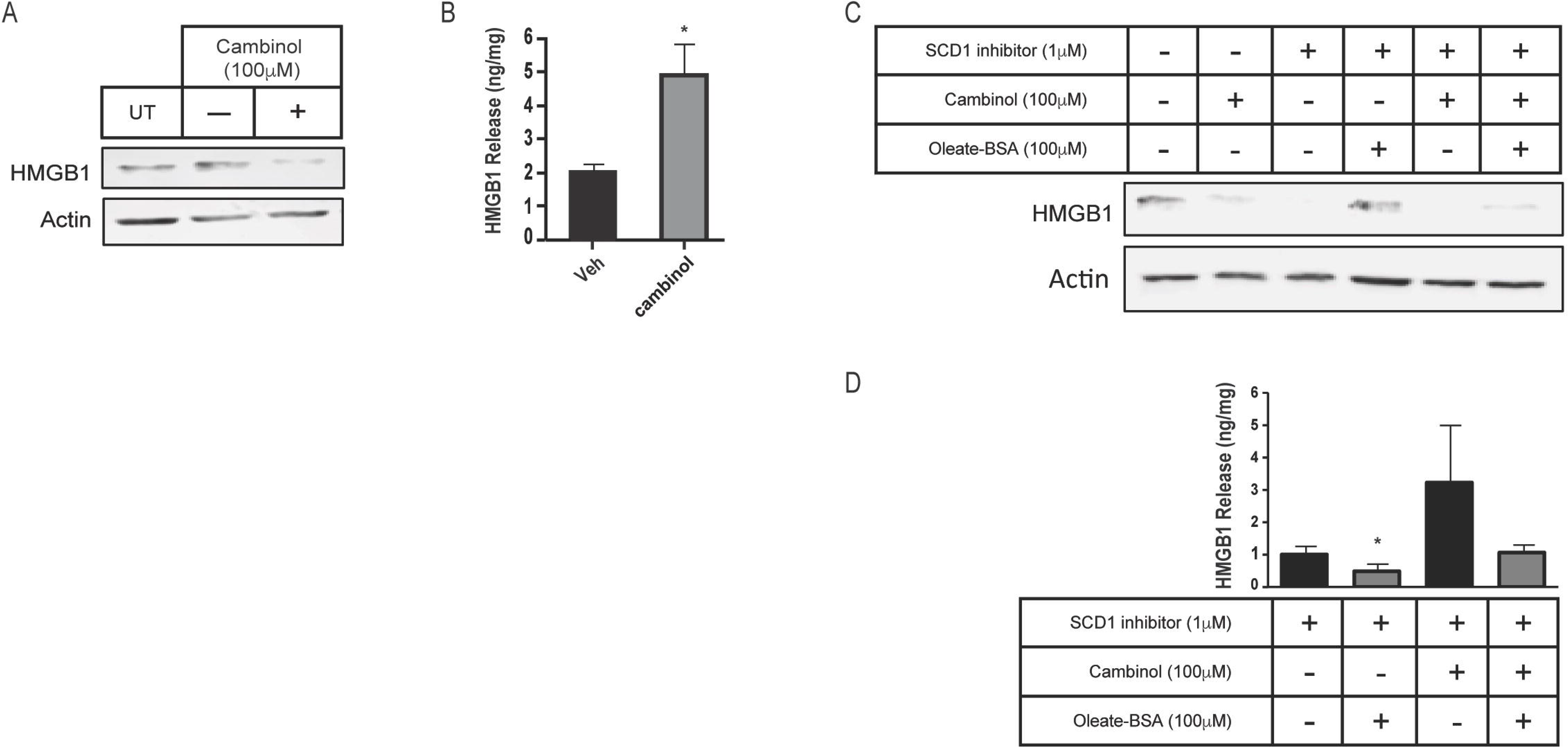
MUFA and SIRT1 cooperate to promote HMGB1 retention in lung cancer cells. A) Immunoblot protein analysis of A549 cells treated for 24 hours with sirtuin 1 and 2 inhibitor, cambinol. B) HMGB1-specific ELISA on extracellular media from A549 cells treated for 24 hours with cambinol. C) Immunoblot protein analysis on A549 cells treated with cambinol in the presence and absence of SCD1 inhibitors and oleate-BSA. D) HMGB1-specific ELISA on extracellular media from A549 cells used in figure 6C. Error bars represent the standard error of the mean (S.E.M.) of three independent experiments. Blots are representative images of three independent experiments.

### MUFA suppresses NF-kB-dependent cytokine release from lung cancer cells

Altering the availability of MUFA has revealed a relationship between fatty acids and protein expression in lung cancer cells. Unsaturated lipids are known to suppress inflammation in cells and tissues. This is associated with UFA reversing ER stress [42]. The transcription factor NF-kB is a significant driver of many inflammatory genes such as TNF alpha, IL-1, and others. To find whether NF-kB was affected by the availability of MUFA, we measured the cytokine response in lung cancer cells in the presence and absence of MUFA. Following delipidation, replenishment of MUFA significantly decreased the amount of DNA-bound nuclear NF-kB p65 (Figure 7A). We next confirmed that the decrease in nuclear NF-kB p65 was associated with changes in cytokine profiles via ELISA. We observed that MUFA decreased the release of NF-KB-driven TNF-a, with and without delipidation (Figure 7B). Interestingly, there was no apparent effect on IL-6 secretion (Figure 7C). The lack of a response in IL-6 may be due to differential regulation, as STAT transcription factors can transcriptionally regulate IL-6. This led us to think that part of the effect MUFA has on lung cancer cells involves regulating inflammatory signaling via HMGB1.

**Figure 7.**
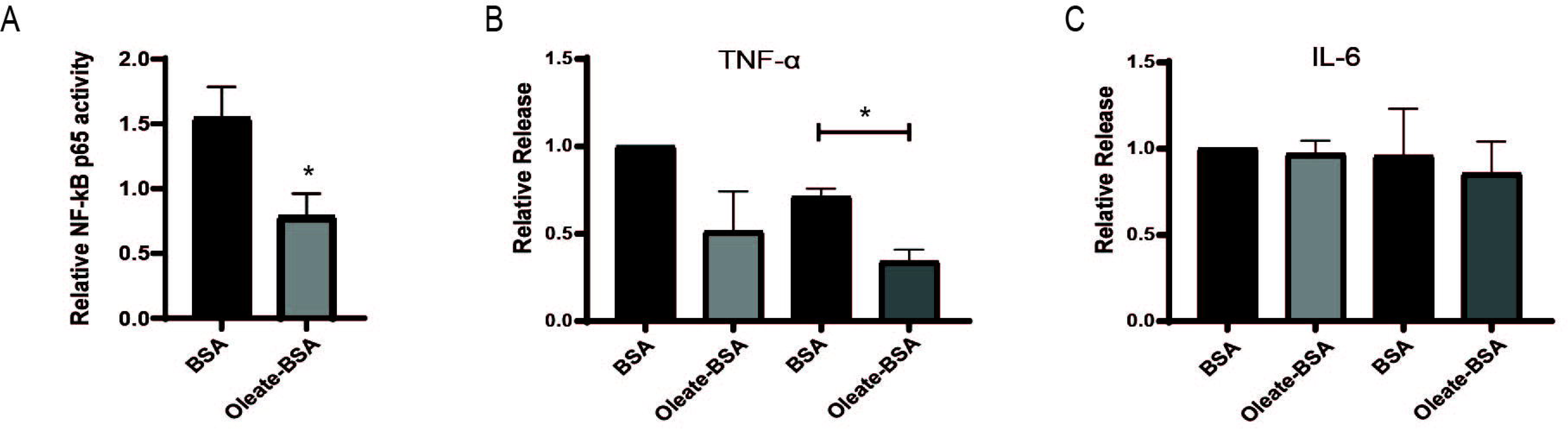
MUFA suppresses NF-kB-dependent cytokine release from NSCLC cells. A) Relative NF-kB p65 DNA binding in A549 cells delipidated for 16 hours and replenished with 100 µM oleate-BSA for 4 hours. B and C) Relative release of inflammatory cytokines from A549 cells treated with 100uM oleate-BSA alone, or with delipidation and replenishment with oleate-BSA. Error bars represent the standard error of the mean (S.E.M.) of three independent experiments. *, indicated a p value<0.05.

### MUFA suppresses inflammatory signaling in monocytes

Cancer cells do not act in a vacuum and often communicate with the surrounding stroma and immune cells to create an immune microenvironment that supports tumor growth. We took conditioned media from delipidated lung cancer cells and placed it on monocytic THP-1 cells to investigate the interaction between lung cancer cells and innate immune cells. Conditioned media from MUFA-treated lung cancer cells suppressed secretion of TNF-a but did not significantly alter other inflammatory markers (MIP-1a and IL-6) (Figure 8A, B). Conversely, conditioned media from MUFA-treated lung cancer cells blunted the release of anti-inflammatory IL-10 from monocytic cells (Figure 8A). This result demonstrates that the availability of MUFA to lung cancer cells may suppress monocyte/macrophage behavior in a manner that influences the tumor microenvironment and subsequent immunogenic behavior.

**Figure 8.**
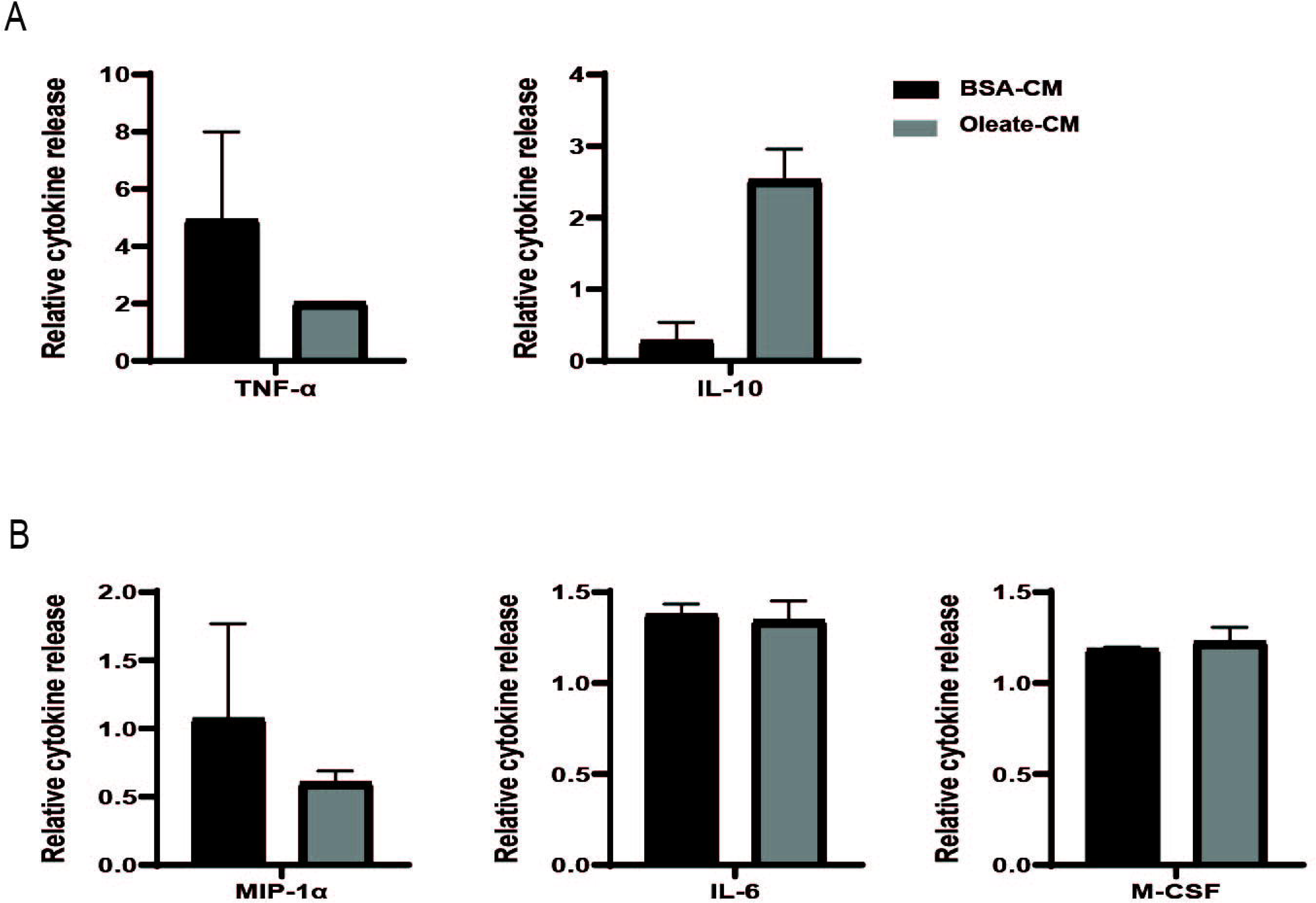
MUFA suppresses conditioned media-induced inflammatory signaling in monocytes. A) Relative release of TNF-alpha and IL-10 from THP-1 monocytes following 24-hour exposure to conditioned from delipidated and oleate-BSA replenished A549 cells. B) Relative release of additional proinflammatory and monocyte-derived cytokines from THP-1 monocytes following 24-hour exposure to delipidated A549 conditioned media. Error bars represent the standard error of the mean (S.E.M.) of three independent experiments. *, indicated a p value<0.05.

### Decreased HMGB1 protein diminishes conditioned media effect on monocytes

HMGB1 in the immune response has a variety of interactions with receptors, including the TLRs and RAGE. The multiple binding partners allow HMGB1 to be involved with various pathogen and damage signaling responses. Therefore, we decided to use a monocyte reporter cell system to identify which pathway was critical to the MUFA-HMGB1 influence on the immune response in lung cancer. We used genetic and pharmacologic methods to inhibit HMGB1 in conditioned media from MUFA-treated lung cells. HMGB1-specific siRNA dramatically decreased in total HMGB1 protein (Figure 9A). siRNA-mediated knockdown was accompanied by a later decrease in HMGB1 released from cells; however, HMBG1 release remained sensitive to adding free fatty acids. Adding MUFA decreased HMGB1 release in siRNA-treated cells, even though HMGB1 levels were already down 50% (Figure 9B). To measure the effect of HMGB1 knockdown in lung cancer cells on monocytic cells, we incubated monocyte reporter (NF-kB and IRF3) cells with HMGB1-depleted lung cancer-conditioned media. MUFA suppressed the stimulation of the IRF3 and NF-kB reporters by conditioned media but was significantly stimulated by the presence of saturated fatty acid (Figure 9C). Interestingly, HMGB1 knockdown significantly suppressed IRF3 and NF-kB activity in all conditions, but NF-kB activity was further reduced with the addition of oleate acid (Figure 9C upper). This suggests that the effect of oleic acid and HMGB1 knockdown are redundant in IRF3 signaling but not for NF-kB.

**Figure 9.**
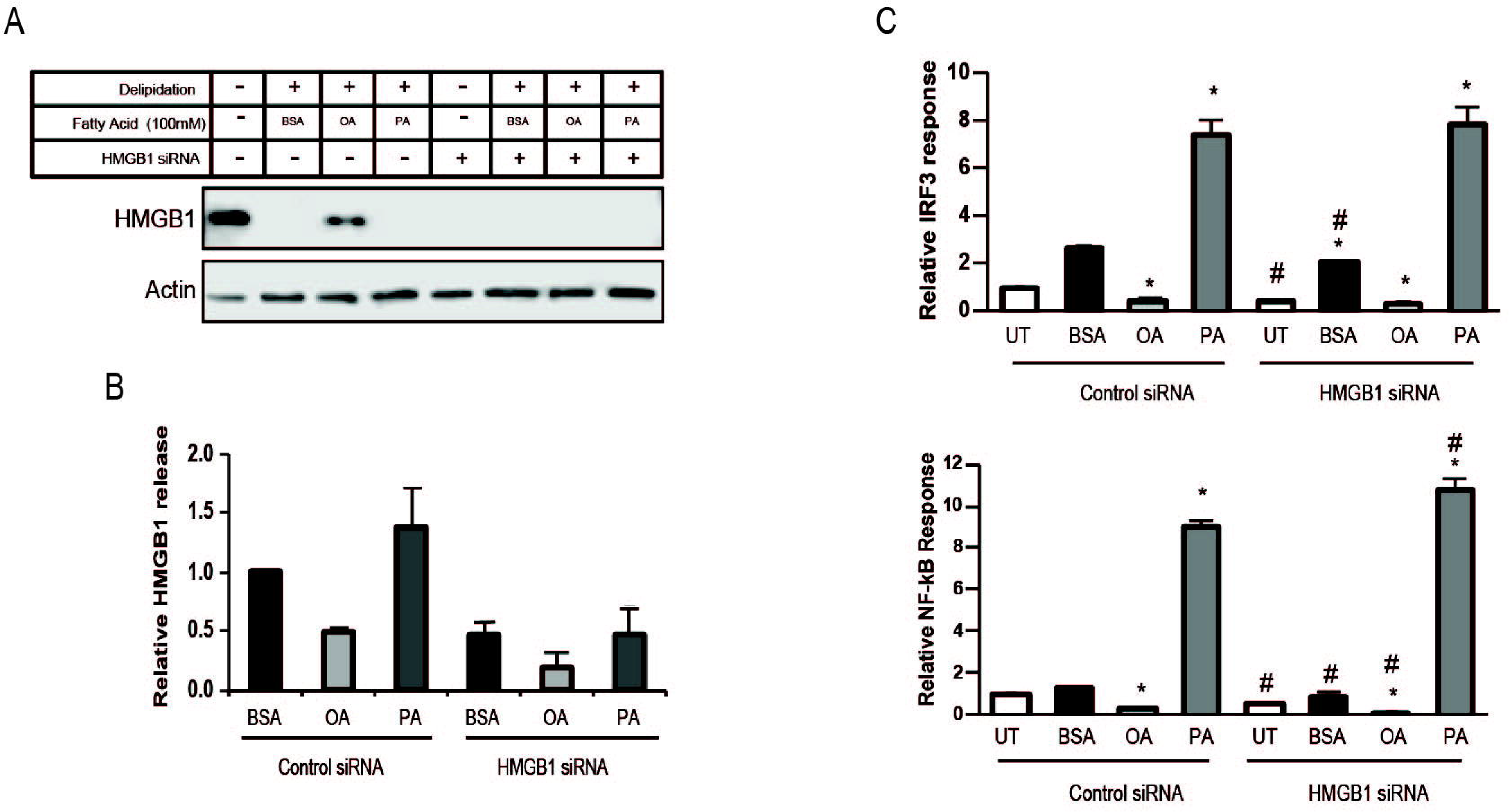
Decreased HMGB1 diminishes impact of A549-cond. media on monocytes. A) Immunoblot protein analysis of delipidated A549 cells replenished with BSA, oleic acid-BSA, or palmitic acid-BSA; 72 hours after transfection with HMGB1-specific siRNA. B) HMGB1-specific ELISA on extracellular media from cells described in Figure 9A. C) THP-1 Dual (IRF3-Luc and NF-kB-SEAP) reporter assay following 24-hour incubation of reporter cells with conditioned media from A549 cells described in Figure 9A. Error bars represent the standard error of the mean (S.E.M.) of 3 independent experiments. *, indicated a p value<0.05.

To confirm the specificity of the effects of HMGB1 knockdown, we next pharmacologically inhibited HMGB1 with the glycyrrhizin. Treatment of cancer cells with glycyrrhizin decreased the activation of both NF-kB and IRF3 reporters; however, a noticeable dose dependence was only present within the IRF3 reporter (Figure 10A, B). To observe the interaction between MUFA- and glycyrrhizin-mediated suppression of the IRF3 reporter, we briefly added MUFA to glycyrrhizin-treated A549 lung cancer cells. Then, we used the conditioned media to stimulate reporter cells. MUFA and glycyrrhizin treatment enhanced the suppressive effect of lung cancer-conditioned media on IRF3 activation compared to MUFA alone (Figure 10C, D). These findings suggested that the presence of HMGB1 and the level of MUFA significantly affect the ability of lung cancer cells to influence the immune function of myeloid-derived cells in the tumor microenvironment.

**Figure 10.**
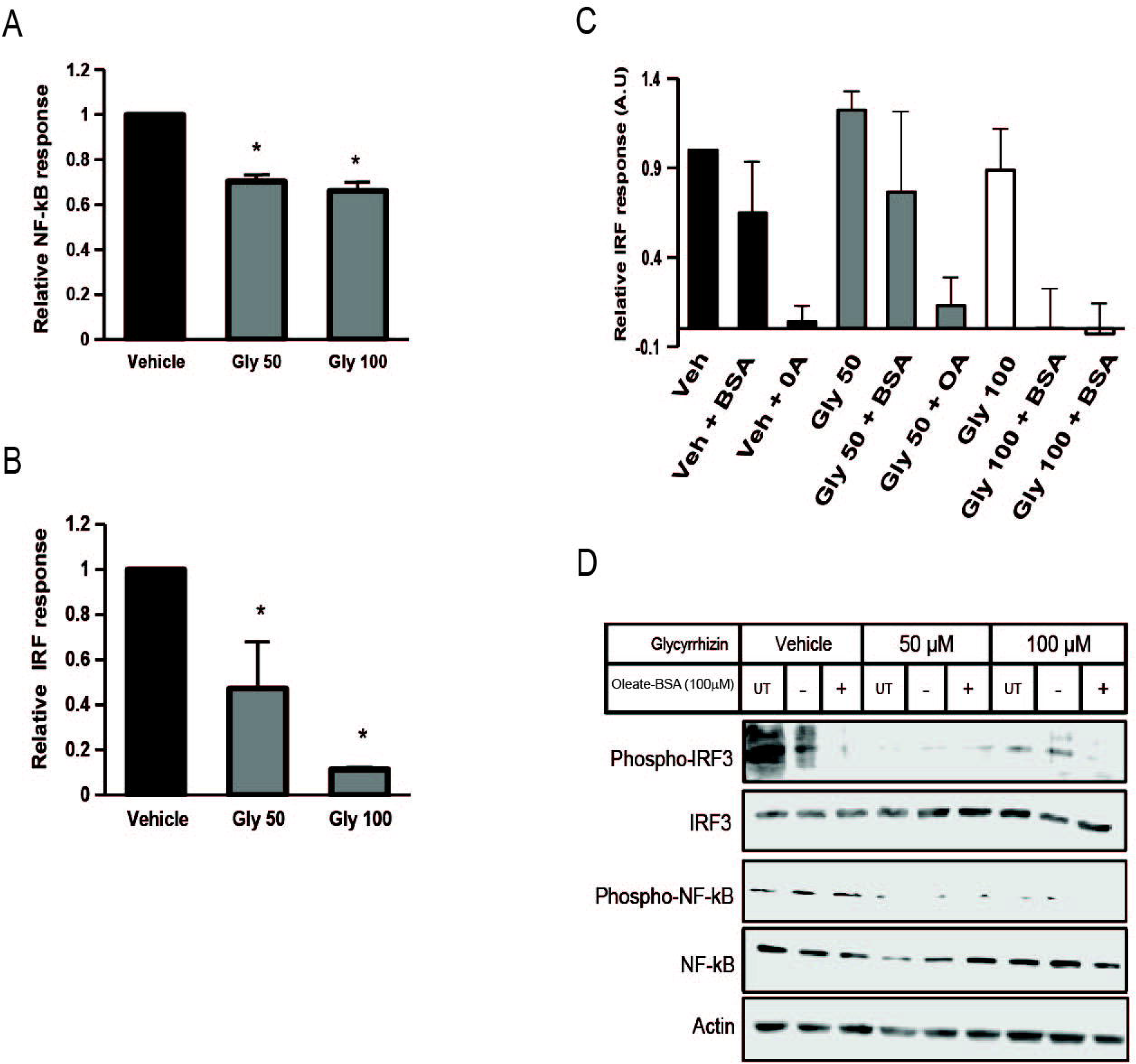
HMGB1 inhibition abrogates IRF3 stimulation in monocytes. A) THP-1 Dual NF-kB reporter analysis following monocyte incubation with conditioned media from A549 cells treated with increasing concentration of HMGB1 inhibitor glycyrrhizin. B) THP-1 Dual IRF3 reporter analysis on the same samples described in Figure 10A. C) THP-1 Dual IRF3 reporter analysis on monocytes incubated with conditioned media from A549 cells treated with glycyrrhizin in presence or absence of oleate-BSA. D) Immunoblot protein analysis of THP-1 Dual cells described in Figure 10C. Error bars represent the standard error of the mean (S.E.M.) of three independent experiments. *, indicated a p value<0.05.

### MUFA increases cellular retention of HMGB1 and decreases PD-L1 expression in lung cancer cells

As we began to view HMGB1 as a MUFA-regulated protein, it became essential to identify its role in the tumor immune landscape. Others have shown that HMGB1 regulates programmed death receptor ligand (PD-L1), a significant target for immune checkpoint therapy. We became interested in finding whether MUFA could also control the expression of PD-L1 via the regulation of HMGB1. To examine the relationship between MUFA and PD-L1, we exposed lung cancer cells to increasing concentrations of MUFA and measured protein expression. We saw that PD-L1 levels diminished as the concentration of MUFA was increased (Figure 11A). Next, we explored how conditioned media from MUFA-treated cancer cells affected PD-L1 expression in immune cells. We exposed monocytes to conditioned media from MUFA-treated A549 lung cancer cells and measured protein expression the following day. Monocytes exposed directly to MUFA had no noticeable changes in PD-L1 protein, but monocytes exposed to MUFA-treated cancer cell-conditioned media dramatically decreased PD-L1 (Figure 11B). The relative decrease in PD-L1 observed in the monocytes supports the idea that MUFA availability affects the ability of cancer cells to influence the immune response in a paracrine-like manner.

**Figure 11.**
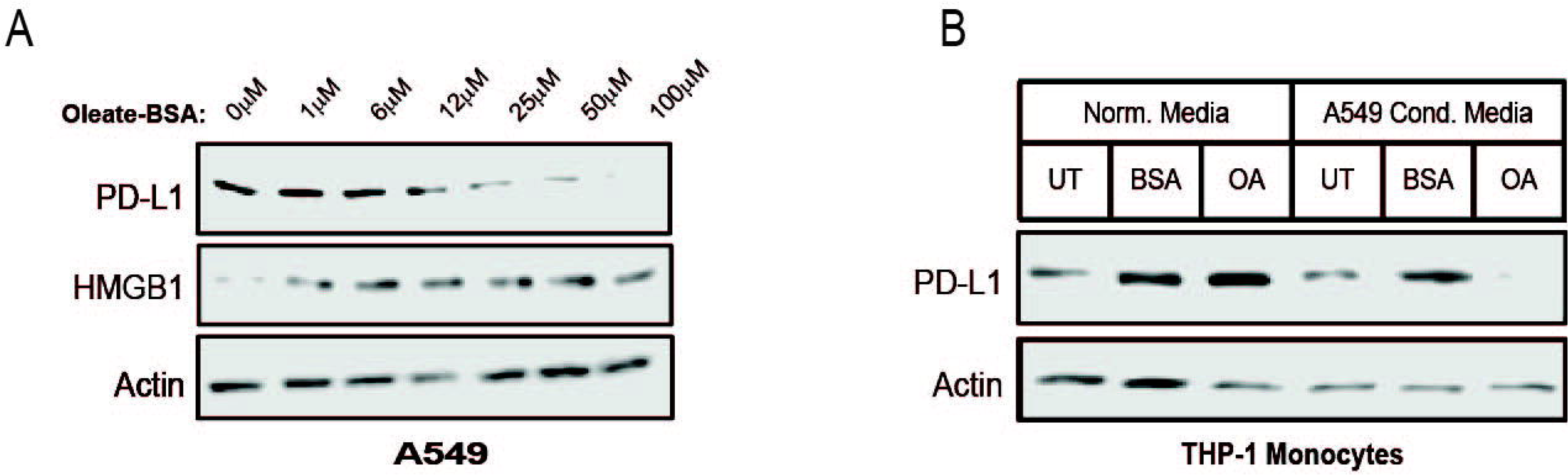
MUFA increases cell-associated HMGB1 and decreases PD-L1 in NSCLC and monocytes. A) Immunoblot protein analysis of HMGB1 and PD-L1 expression in A549 cells treated for 4 hours with increasing concentrations of oleate-BSA. B) Immunoblot protein analysis of PD-L1 expression in THP-1 monocytes following 24-hour incubation in presence or absence of delipidated A549 conditioned media. Blots are representative images of three independent experiments.

### HMGB1 is negatively correlated with PD-L1 expression in NSCLC patients

Our *in vitro* studies suggest that the availability of MUFA would affect the expression of PD-L1 in patient tumors. To investigate this hypothesis, we analyzed the expression of lipogenic genes, body mass index, and CD274 (PD-L1) levels from published multi-omics lung adenocarcinoma datasets (Figure 12A). Multivariate analysis revealed several significant relationships. One of the major findings from this dataset was a significant negative correlation seen between HMGB1 and CD274 (PD-L1) protein in both male and female patients (Figure 12A, B, C). This was due to the absence of HMGB1 within the cell following its secretion and binding to TLR or RAGE receptors on nearby cells. We next observed a positive correlation between SCD1 and PD-L1 protein expression; however, significance was only seen in the female patients (Figure 12B).

**Figure 12.**
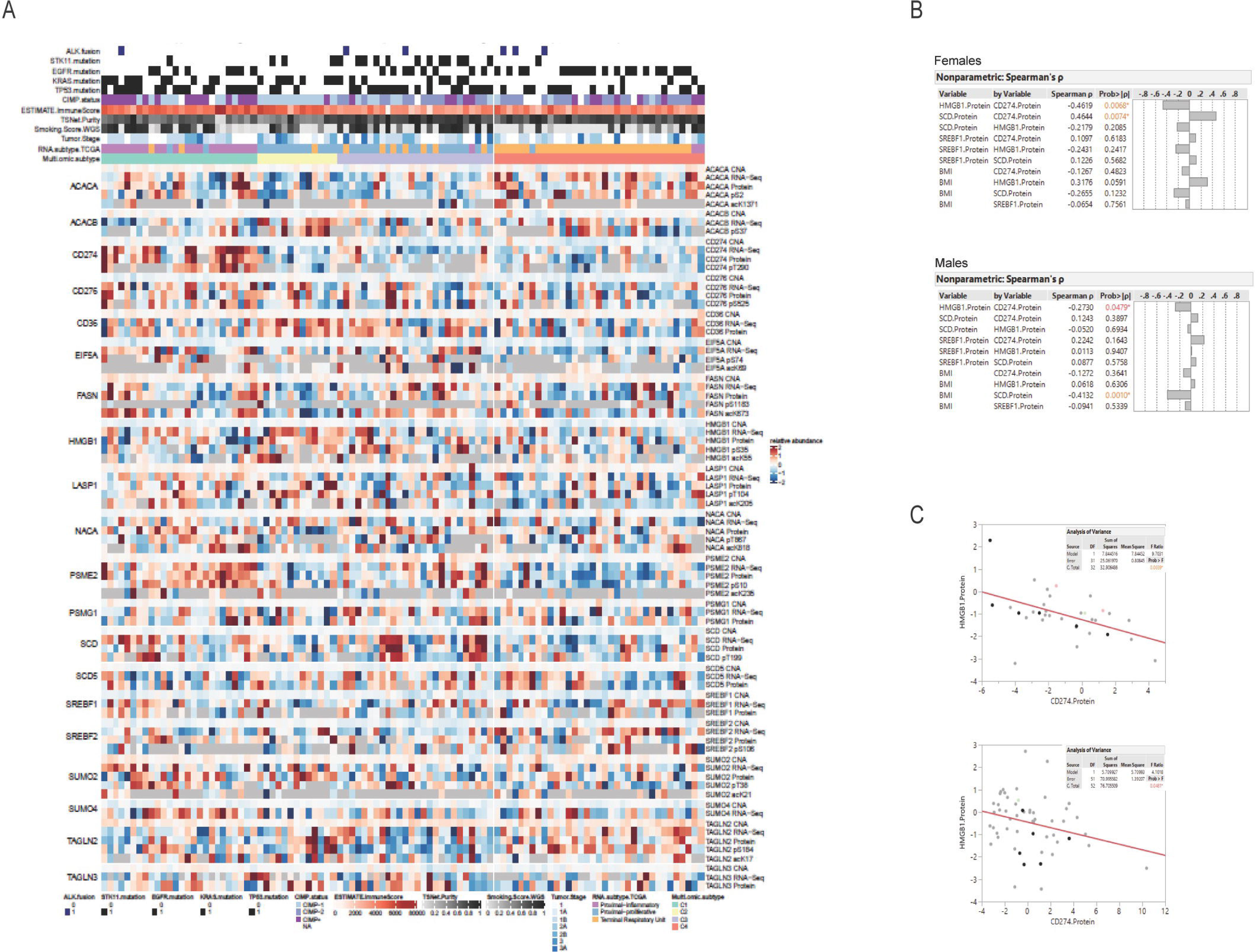
HMGB1 is negatively correlated with PD-L1 in NSCLC patients. Heatmap of multi-omic data from lung adenocarcinoma patient published dataset examining expression HMGB1, SCD1, SREBF1, and PD-L1 (CD274). B) Spearman’s correlation analysis among the select proteins from Figure 12A and body mass index in female and male lung adenocarcinoma patients. C) Pearson correlative analysis between expression of HMGB1 protein and PD-L1 (CD274) in female and male lung adenocarcinoma patients. *, indicates statistical significance and p-value <0.05. n = 111 lung cancer patients.

Although that relationship contradicts our first supposition that SCD1(MUFA) was negatively related to CD274 (PD-L1), it may also indicate a more complicated relationship than we had previously thought. We also observed an unimpressive negative relationship between HMGB1 and SCD1; additional analysis across cancer sub-types further revealed more dramatic statistical differences between HMGB1 and SCD1 within the "proximal inflammatory" and "terminal respiratory unit" subtypes (not shown) [43]. Lastly, as there exists some debate around the role of obesity or elevated BMI and patient prognosis and response to treatment, we also wanted to analyze the relationship between BMI and proteins of interest. Interestingly, we saw a negative relationship between SCD1 and BMI, especially in male patients. This suggests that individuals with elevated BMI also have lower SCD1protein, thus less MUFA available to prevent HMGB1 release, which could drive immune suppression.

To confirm the observed correlation between HMGB1 and PD-L1 in the above dataset, we collected patient serum and tissues to analyze the level of secreted HMGB1 in patients with PD-L1-positive tumors. From the 14 patients enrolled, we obtained serum, tissue, and pathology reports for only five. We measured the amount of HMGB1 in the patient’s serum using a human HMGB1 ELISA. Patients with tumors that were negative for PD-L1 based on immunohistochemical analysis tended to have less HMGB1 in their blood when compared to patients with PD-L1-positive tumors (Figure 13). Acquisition and analysis of more patient tissue and blood from multiple sites are in progress to improve the statistical power of our observations. Withstanding the small sample size, the data we have generated suggests that the relationship between HMGB1, SCD, and PD-L1 could help clinicians stratify patients into groups based on the potential to predict the immunological sensitivity of tumors to current and future immunotherapies.

**Figure 13.**
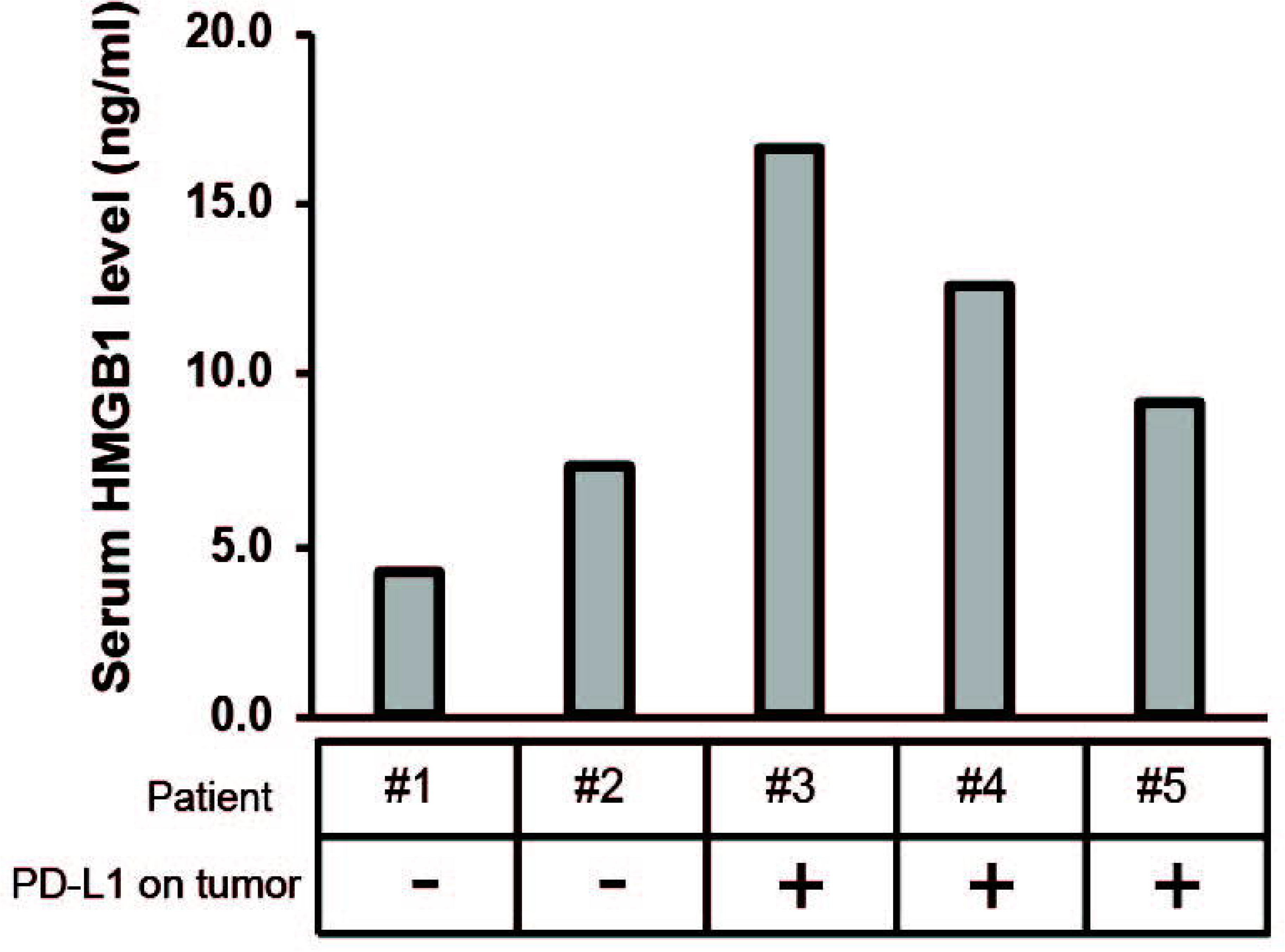
Serum HMGB1 is elevated in patients with PD-L1 tumor staining. HMGB1-specific ELISA on serum samples from lung adenocarcinoma patients, compared to histological detection of tumor-associated PD-L1. Any signal was considered positive for PD-L1 expression using the clinically validated Ventana assay.

## Discussion

In this study, we have uncovered a complex interplay between lipid metabolism, the release of HMGB1, and immune modulation within the context of NSCLC Our findings highlight the critical role of SCD1 and MUFA in regulating essential elements of the tumor microenvironment, shedding light on potential therapeutic strategies and avenues for further investigation.

Our results showed that a significant decrease in the expression of SCD1 exists in NSCLC tumors compared to normal tissues. This finding underscores the importance of lipid metabolism in the context of lung cancer, a concept supported by earlier work from our laboratory and others. The intricacies of de novo synthesis and uptake of MUFA are likely to play pivotal roles in developing and supporting lung tumors. Although the influence of SCD1 expression on patient survival was primarily clear in specific population quartiles, indicating an outlier effect, the observed disparity between SCD1 expression in normal and malignant tissues cannot be overlooked. These results are evidence of the impact of SCD1 and lipid metabolism on tumor progression.

Our results also showed that lipid metabolism plays a significant role in HMGB1 and the immune response to lung cancer. Inhibiting HMGB1 decreased the expression of distinct immune response-related genes, particularly those regulated by NF-kB signaling, affirming that HMGB1 is a critical mediator in the communication between lung cancer cells and the immune system.

Notably, our study unveiled a connection between MUFA availability and the regulation of PD-L1 expression in lung cancer cells, a pivotal target for immunotherapy. PD-L1 expression decreased as the concentration of MUFA increased, showing a potential avenue for modulating the immune response. Furthermore, our analysis of patient data suggested a complex relationship between HMGB1, SCD1, and PD-L1, potentially influencing the immunological sensitivity of tumors to current and future immunotherapies.

In conclusion, this research reveals a multifaceted relationship between lipid metabolism, HMGB1, and immune modulation within the context of NSCLC. These findings open doors to further investigations and the development of innovative therapeutic strategies that target key elements of this intricate interplay. Understanding the regulation of lipid metabolism and immune responses in lung cancer is crucial for developing more effective treatments and personalized approaches for patients. As we continue to uncover the complexities of these interactions, we hope to contribute to the ever-advancing field of cancer research and the eventual improvement of patient outcomes.

## Materials and Methods

### Cell Culture

A549, HCC827, H1299, H23 non-small cell lung carcinoma cells were purchased from ATCC. A549 and H1299 cells were cultured and maintained in low glucose DMEM (Corning cat #10-014-CV) supplemented with 5% FBS, 50 U /ml penicillin, 50 μg ml/ml streptomycin. Hcc827 and H23 cells were cultured and maintained in low glucose DMEM supplemented with 10% FBS, 50 U /ml penicillin, 50 μg ml/ml streptomycin. THP1-Dual cells (InvivoGen) were cultured and maintained in RPMI 1640 (Corning) supplemented with 10% FBS, 50 U /ml penicillin, 50 μg ml/ml streptomycin, 100 μg/ml normocin, and 2 mM L-glutamine.100 μg/ml of zeocin and 10 μg/ml of blasticidin were added to growth medium every other passage to retain the dual reporters.

### Lipid deprivation

Lung cancer cells were plated in complete DMEM as described above at appropriate cell density for specific experimental conditions overnight. The following day, all culture media was removed, and cells were rinsed one time with sterile phosphate buffered saline. Cells were replenished with delipidation media consisting of DMEM supplemented with 100 units/ml penicillin, and 100 µg/ml streptomycin with 5% delipidated fetal calf serum (DFCS), and 1 µM SCD-1 inhibitor (A939572, Cayman Chemicals). Cells were incubated in delipidation media overnight and assayed for response to lipids the following day. The lipid replenishment was carried by adding the desired amount of specified fatty acid for 4-6 hours.

### Cell harvest and lysis

Cells were harvested by scraping the cells and media into conical tubes and placing them on ice. Following harvest, the cell suspensions were spun down at room temperature for 5 min at 800 × g. The supernatant was aspirated, and the cells were washed twice with PBS. Cells were then lysed with 100 µl of seize2 lysis buffer (25 mM Tris-HCl pH 7.2, 0.15 M NaCl, 1% NP-40, and 1x cOmplete EDTA-free protease inhibitor cocktail (Roche cat no. 4693132001). The lysate was passed through a 22 G needle ten times while on ice followed by rotation at 4°C for 15 min. The lysate was pelleted at 21,000 × g for 10 min and the supernatant was collected. Protein concentrations were found using the BCA Protein Assay kit (Thermo Fisher Scientific) and read at 562 nm. Sample lysate was mixed with loading buffer and then boiled for 5 min and loaded onto 10% SDS-PAGE gel at 10 or 20 µg total protein per well.

### Immunoblot analysis

Following overnight treatment with fatty acids cells were harvested and lysed as previously described. A total of 10 µg of protein was then separated by 10% SDS-PAGE. After gel electrophoresis was completed, the proteins were transferred to a 0.2-micron polyvinylidene difluoride (PVDF) membrane. All immunoblot equipment and reagents used were from Bio-Rad. The membrane was blocked with a 5% non-fat milk/PBST solution for 30 min. The membrane was incubated with primary antibodies at 4°C overnight while rocking. The following day the membrane was washed three times with PBST at 5 minutes per wash. Then 5% non-fat milk/PBST containing HRP-conjugated secondary was added to the membrane. β-actin was used as the loading control for all immunoblots (primary antibodies: Sigma, 1:2,500 dilution; secondary antibody. Protein bands were detected by using 1 ml of SuperSignal ECL from Pierce and visualizing band intensity using Li-Cor C-Digit imaging system.

### Cell Viability

For the CCK-8 assay, A549 cells with or without SCD1 inhibition were seeded at a density of 1000 cells/well in 96-well plates for 24 hours. Subsequently, 10 μl of CCK-8 solution (Abcam, cat no. ab228554) was added to each well, and the plates were incubated at 37 °C for 2 h. Finally, the absorbance was measured at 450 nm.

### Plasmid transfection

A549 cells were seeded in 60 mm plates then transfected with 2 µg of N-terminal tagged GFP-HMGB1 DNA plasmid DNA (Sino biological, cat no. HG10326-ANG). In brief, 250,000 cells were plated in 60 mm dishes the night prior to transfection and placed at 37°C. Transfection were done using X-tremeGENE HP DNA transfection reagent (Roche, cat no. 6366244001). Each dish was transfected with 2 µg of plasmid DNA per manufacturer’s instructions. The GFP signal was confirmed using a CellDrop Automated Fluorescent Cell Counter (DeNovix) to confirm protein expression prior to imaging on the EVOS M5000 fluorescent imaging system (ThermoFisher, cat no. AMF5000). Cells were incubated 24 h before harvest to analyze protein and mRNA expression.

### Fluorescent Microscopy

Following treatment of HMGB1-GFP transfected A549 cells with oleate-BSA cells were stained with Nile Red. In brief, stock solution of Nile Red (1mg/ml in DMSO) was diluted to 1mg/ mL. Cells were washed twice with sterile PBS and fixed with 3.7% formaldehyde for 15 minutes at room temperature. Formaldehyde was wash away with two PBS washes before transferring coverslips to microscope slide for imaging on EVOS M500 imaging system using 20x magnification.

### RNA interference

150,000 cells were plated in 60 mm dishes overnight at 37°C. Following day transfection mixes were made for each experimental permutation. SiGENOME SMARTpool siRNA for HMGB1 (M-018981-01-0005) and control non-targeting siRNA pool #1 (D-001206-13-05) were transfected 100 nM per dish. Lipofectamine RNAiMax (Invitrogen Cat: 13778030) was mixed with appropriate volume of OptiMEM (Gibco) in one tube, while in a separate tube the appropriate amount of each siRNA was added to appropriate volume of OptiMEM (Gibco, cat no. 31985070) and mixed. Individual tubes were incubated at room temperature for 5 min. The two solutions were combined at equal volume, mixed, and incubated at room temperature for 20 min. Plates containing cells, were washed, and replenished with 1.6 ml of OptiMEM, and placed back at 37°C until transfection complexes were ready. Then 400 µl of transfection mix was added to each dish, and incubated at 37°C for 4 h, at which cells would be washed once with complete media and 4 ml of complete media would be added to cells. Cells then incubated at 37°C for 48 h.

### Reverse Transcriptase Quantitative polymerase chain reaction

24-h post transfection with siRNAs as described above. Total RNA was obtained from cells by Qiazol extraction and RNeasy purification (Qiagen, cat no. 74104). HMGB1 and SCD1 mRNA levels were determined using RT-qPCR using a Rotor-gene Q PCR machine. The data were analyzed using the 2−ΔΔCq method and the Actin mRNA was used as an endogenous control as previously described [44].

### Lucia luciferase and SEAP assays

QUANTI-Luc luciferase reagent (InvivoGen, rep-qlc4lg5) was used to measure IRF activity through Luciferase luminescence following the manufacturer’s protocol. To measure IRF, 20 μl of cell culture supernatant was transferred to 96-well opaque plate and luminescence was read using a Biotek Synergy spectrophotometer after addition of 50 μl of luciferase reagent. NF-kB activity was measured by combining 20 μL of supernatant with180 μL QUANTI-Blue SEAP detection reagent (InvivoGen, cat no. rep-qbs) in 96-well assay plate. The samples were then incubated at 37 °C for 1 h and absorbance was measured at 650 nm.

### Statistical analysis

All experiments were performed in triplicate (n=3), unless otherwise indicated. An unpaired two-tailed Student’s t-test with two degrees of freedom was used to compare means of the three replicate experiments between treatments using either GraphPad Prism or MS Excel. Where appropriate, the Bonferroni correction was applied to t-tests. P<0.05 was considered to indicate a statistically significant difference. Association between OS and key proteins was determined by univariable Cox proportional Hazard models. Associations between key proteins of interest were determined by multivariate pairwise analysis using Spearman ranked correlations for each pair of set of variables (P<0.05 implies a statistically significant marginal association at the 0.05 alpha level). Multivariate analysis was done using proportion of pairwise correlations in JMP software by SAS. Far publicly available patient data, the method for differential analysis is one-way ANOVA, using disease state (Tumor or Normal) as variable for calculating differential expression. The expression data are first log 2(TPM+1) transformed for differential analysis and the log 2FC is defined as median(Tumor) - median(Normal). Genes with higher |log 2FC| values and lower q values than pre-set thresholds are considered differentially expressed genes.

## Data availability

The authors confirm that data supporting the findings of this study are available within the article and the publicly available data sources that have been mentioned in the manuscript. Proteomic analysis of CPTAC LUAD was performed using the CPTAC LUAD Data Viewer (https://prot-shiny-vm.broadinstitute.org:3838/CPTAC-LUAD2020/). Differential expression of SCD1, HMGB1 genes from the TCGA LUAD dataset was analyzed on Gene Expression Profiling Interactive Analysis (GEPIA2) (http://gepia2.cancer-pku.cn).

## Author Contributions

GES: Study concept and design. GES: Acquisition of funding. GES, BCS, NMA, ZB, KR: Acquisition of data. GES, BCS, NMA, ZB: Analysis and interpretation of data. GES, BCS, NMA, ZB, KR: Drafting of the manuscript. GES, BCS, NMA, ZB, KR: Critical revision of the manuscript for important intellectual content.

## Acknowledgements

The authors thank the present and the past members of the Simmons lab for supporting the study. Proteomic analysis non-small cell carcinoma cell lines were done by the Proteomics and Metabolomics Facility at the Cornell Institute of Biotechnology (RRID:SCR_021743). The results shown here are in whole or part based upon data generated by The Cancer Genome Atlas (TCGA) Research Network: https://www.cancer.gov/tcga.

